# Leaf shape is a predictor of fruit quality and cultivar performance in tomato

**DOI:** 10.1101/584466

**Authors:** Steven D. Rowland, Kristina Zumstein, Hokuto Nakayama, Zizhang Cheng, Amber M. Flores, Daniel H. Chitwood, Julin N. Maloof, Neelima R. Sinha

## Abstract

- Commercial tomato (*Solanum lycopersicum*) is one of the most widely grown vegetable crops worldwide. Heirloom tomatoes retain extensive genetic diversity and a considerable range of fruit quality and leaf morphological traits.
- Here the role of leaf morphology was investigated for its impact on fruit quality. Heirloom cultivars were grown in field conditions and BRIX by Yield (BY) and other traits measured over a fourteen-week period. The complex relationships among these morphological and physiological traits were evaluated using PLS-Path Modeling, and a consensus model developed.
- Photosynthesis contributed strongly to vegetative biomass and sugar content of fruits but had a negative impact on yield. Conversely leaf shape, specifically rounder leaves, had a strong positive impact on both fruit sugar content and yield. Cultivars such as Stupice and Glacier, with very round leaves, had the highest performance in both fruit sugar and yield. Our model accurately predicted BY for two commercial cultivars using leaf shape data as input.
- This study revealed the importance of leaf shape to fruit quality in tomato, with rounder leaves having significantly improved fruit quality. This correlation was maintained across a range of diverse genetic backgrounds and shows the importance of leaf morphology in tomato crop improvement.

## Introduction

The rise of agriculture around 7000 BC ensured a stable food supply, allowing human civilizations to develop and populations to grow (Barker, 2006). We have now come full circle with population growth fast outstripping the capacity of current agricultural output. Added to the challenge of growing populations is climate unpredictability, with drought and temperature increases leading to decreased crop yield (Matiu *et al.*, 2017). Tomato (*Solanum lycopersicum*) is by far the most widely grown vegetable crop worldwide (Bauchet and Causse, 2012). The narrow genetic base of most crops, combined with selection for performance under optimal conditions, has reduced the genetic variability in environmental stress responses, and the modern cultivars of tomato are no exception (Bai and Lindhout, 2007; Bauchet and Causse, 2012; Bergougnoux, 2013). The wild relatives of tomato have the genetic ability to adapt to extreme habitats, and many heirloom cultivars also retain this ability due to outcrossing with wild species, and less breeding selection for commercially valuable traits (Lin *et al.*, 2014; RodrÍGuez-Burruezo *et al.*, 2015). Heirloom tomatoes are varieties which have been passed down through multiple generations of a family (traditional), or open pollinated for more than 50 years (commercial varieties; Tomato Fest, https://www.tomatofest.com/what-is-heirloom-tomato.html). Improvement in tomato has focused on flowering, fruit traits, and disease resistance, while other potential impacts of vegetative organs on yield and fruit quality are relatively ignored (Grandillo *et al.*, 1999; Passam *et al.*, 2007; Bergougnoux, 2013; Bauchet and Causse, 2012).

A previous study by Chitwood and coworkers (2013) linked leaflet shape in tomato to fruit sugar content. It was shown that plants with rounder leaflets also had increased fruit sugar content (Chitwood *et al.*, 2013). Because leaves are the primary site of photosynthesis, it is possible that leaf shape changes may impact photosynthetic capacity and therefore result in different levels of sugar content (BRIX) and yield in fruits. In addition to photosynthesis, sugar transport, and distribution to sinks are other potential sites of regulation in leaf function as source tissue. While sugar transport in plants is well described, distribution among different sink tissues is not fully understood (Lemoine *et al.*, 2013).

To elucidate the relative importance of leaf shape to tomato fruit traits, we analyzed a set of eighteen heirloom tomato cultivars with varied yield and fruit quality, photosynthetic capacity, leaflet shape, and other vegetative traits. Additionally, we performed WGS analysis to determine the genetic relationship among a subset of these lines as they have diverse genetic backgrounds and breeding histories. We found that in this selection of heirlooms, leaflet shape was strongly correlated to overall fruit quality assessed as a composite measure of BRIX and Yield (BY), with rounder leaflets positively correlated with higher BY values. Photosynthesis, while having a strong correlation with vegetative biomass and fruit BRIX, had a negative correlation with yield. Based on our analysis, leaf shape seems to play an important role in the distribution of photoassimilates.

## Materials and Methods

### Plant Material and Growth Conditions

Eighteen heirloom tomato varieties identified as having a range of tomato types including cherry and beefsteak tomatoes, and several intermediate types, were analyzed. These tomato varieties also differed in fruit production timing from early to late, and the type of leaf morphology. They were initially selected based on leaf morphology as described in Tatiana’s TOMATObase and The Heirloom Tomato (http://tatianastomatobase.com/wiki/Main_Page; Goldman, 2008).

Tomato seeds were soaked in 50% bleach for 10 min, rinsed 3-5 times with water, and placed on water dampened Phytatrays (Sigma Aldrich). Seeds were incubated in the dark at room temperature for 3 days then transferred to a growth chamber set at 25°C with 16:8 photoperiod for 6 days. After approximately 6 days, seedlings had expanded cotyledons. These were then transplanted to 72-cell Seedling Propagation trays (TO Plastics) and grown in the chamber for 7 days watered with nutrient water. At 15 days old, seedlings were transferred to the greenhouse (end of April). After 2 weeks, seedlings were transferred to the lath house (mid-May). Both in the greenhouse and lath house, plants were watered from the top to encourage hardening. Lath house watering used only RO water to encourage hardy, compact plantlets with strong root plugs for hand or mechanical transplanting in the field. In both the 2014 and 2015 seasons plants were laid out in a randomized block design and were planted (late May) and grown in soil, with furrow irrigation once weekly.

### Gas Exchange and Intercepted PAR Measurements

Gas exchange measurements were made in the field on attached leaves after the plants had recovered from transplanting. Measurements were made weekly from week 10 through week 15 (vegetative growth), week 17 (initiation of flowering), and 18 – 21 (fruiting stages), on approximately 60 plants each week. Measurements were made on leaves from the upper and lower portion of the plants to eliminate positional bias within the plant. The A, gH_2_0, transpiration, and ɸPS2 of a 6 cm^2^ area of the leaflet was measured using the LI-6400 XT infra-red gas exchange system (LI-COR, Lincoln, NE), and a fluorescence head (6400-40; LI-COR, Lincoln, NE). The chamber was positioned on terminal leaflets such that the mid-vein was not within the measured area. Light within the chamber was provided by the fluorescence head at 1500 µmol m^−2^ s^−1^ PAR, and the chamber air flow volume was 400 µmols s^−1^ with the chamber atmosphere mixed by a fan. CO_2_ concentration within the chamber was set at 400 µmols mol^−1^ (average atmospheric concentration). Humidity, leaf, and chamber temperature were allowed to adjust to ambient conditions, however the chamber block temperature was not allowed to exceed 36°C. Measured leaflets were allowed to equilibrate for 2 to 3 minutes before measurements were taken, allowing sufficient time for photosynthetic rates to stabilize with only marginal variation.

The amount of intercepted PAR (Photosynthetically Active Radiation; PARi) was measured in four orientations per plant and an average PARi calculated. PARi was measured by placing a Line Quantum Sensor (LQA-2857; LI-COR, Lincoln, NE) onto a base made from ¼” PVC piping, and a Quantum Sensor (LI-190R; LI-COR, Lincoln, NE) approximately 1 meter above the plant on the PVC rig. Measurements from both sensors were taken simultaneously for each sample using a Light Sensor Logger (LI-1500; LI-COR, Lincoln, NE). This allowed variation in overall light levels such as cloud movement to be measured and accounted for in the total PARi.

### Harvest Measurements

After gas exchange measurements, three plants per genotype were destructively harvested each week. The final yield (weight of all fruit) and fresh vegetative weight of each plant harvested, was measured using a hanging scale (TL 440; American Weigh Scale, Inc., Norcross, GA) in the field. At least five leaves were collected from each plant for later analysis, and approximately nine fruit were collected for BRIX measurements. Fresh weight was used due to the large number of plants and measurements being done *in situ* in the field setting. All measurements were made in kg. To measure the BRIX value of the tomatoes, the collected fruit was taken to the lab where the juice was collected and measured on a refractometer (HI 96801 Refractometer, Hanna Instruments Woonsocket, RI). The yield and BRIX for each plant were multiplied together to get the BRIX x Yield Index (BY), which gives an overall fruit quality measure, accounting for variations and extreme values in either measurement.

### Leaf Morphology Analysis

Leaf complexity and leaflet shapes were analyzed for leaves collected from the field. The leaf complexity measures included all leaflets present on the leaf. After complexity was obtained the primary leaflets were removed and used for imaging and analysis of shape and size. The intercalary and secondary/tertiary leaflets were not used for shape analysis due to their smaller size and irregular shapes. Leaflets were imaged using an Olympus camera (Olympus SP-500UZ) at a fixed distance and the images then processed in ImageJ (Schneider *et al.*, 2012). The images were cropped to individual leaflets maintaining the exact pixel ratio of the original image, and then cropped again to only include the single leaflet using a custom Java script written for FIJI (Schindelin *et al.*, 2012). At the same time these single leaflet images were converted to a binary image so that the leaflet was black on a white background, and the image smoothed to allow for the exclusion of any particulates in the image. The binary images were then processed in R using MOMOCS (Bonhomme *et al.*, 2014), a shape analysis package. Leaflet images were imported and then aligned along their axes so that all images faced the same direction. They were then processed using elliptical Fourier (eFourier) analysis based on the calculated number of harmonics from the MOMOCS package. PCA analysis was performed on the resulting eFourier analysis and the Principal Components (PCs) used for subsequent analysis. Traditional shape measures such as leaflet area, circularity, solidity, and roundness were made with the area measurement based on pixel density. These measures were compared to the PCs to determine the characteristics captured by each PC. The PC values were used for all subsequent leaflet shape and size analyses. Total leaf area for each plant was measured by imaging the whole plant and a 4 cm^2^ red square and then processed in the Easy Leaf Area software (Easlon and Bloom, 2014; Fig. 2b).

### Leaflet Sugars

Five plants per line were used to analyze leaflet sugar content. The plants were grown under the same conditions as field plants with the following exceptions. Plants remained in the greenhouse after transfer to one-gallon pots. All plants were watered with nutrient solution and grown until mature leaves could be sampled. Using a hole punch a disc with an area of 0.28 cm^2^ was taken from a leaflet from each plant. Sugar was extracted from the discs using a modified extraction method from the Ainsworth lab (Bishop *et al.*, 2018). Leaf discs were placed in 2mM HEPES (Affymetrix Inc.) in 80% EtOH (Sigma-Aldrich) and heated to 80°C for 20 minutes and the liquid collected and stored at −20°C. The entire process was repeated twice. They were then placed in 2mM HEPES in 50% EtOH and heated, collecting the liquid and storing at −20°C followed by another 2mM HEPES in 80% treatment. The collected liquid was then used to measure the amount of sugar present per area of disc.

To measure leaf sugar content a working solution of 100mM HEPES (Affymetrix Inc.), 6.3mM MgCl_2_ (Sigma-Aldrich), and 3mM ATP (Sigma-Aldrich) and NADP (Sigma-Aldrich) at pH 7 was prepared. From the working solution an assay buffer was made adding 50U of Glucose-6-Phosphate Dehydrogenase (G6PDH; Sigma-Aldrich), and 295µL or 280µL of the working solution was added to a 96 well plate (Costar) for sucrose standards or samples respectively. Standards were added at a 60-fold dilution and samples were added at a 15-fold dilution. Then 0.5U of Hexokinase (Sigma-Aldrich), 0.21U of Phosphoglucoisomerase (PGI; Sigma-Aldrich), and 20U of Invertase (Sigma-Aldrich) were added to each well and the plates allowed to sit overnight to reach equilibrium. The plates were measured on a UV-Spectrometer (Molecular Devices SPECTRAmax 340) at 340 nm, followed by analysis in JMP (JMP Pro 14.0.0, 2018 SAS Institute Inc.). The total sugar content was measured using this method.

### Statistical Analysis

All statistical analyses were performed using JMP (JMP Pro 14.0.0, 2018 SAS Institute Inc.) software. To determine statistical significance, measurements were modeled using General Linear Regression Model (GLM) and tested by a One-Way ANOVA followed by Tukeys-HSD, if necessary. This modeled data for all measured values was compiled into a table and used to create a model using the Partial Least Squares-Path Modeling (PLS-PM) in SmartPLS3.0 (Ringle *et al.*, 2015). Modeled data was used for the statistical analyses as many measurement types varied in number of data points, and therefore a set of generated predicted values of equal size was used to make an equal data matrix.

PLS-PM was used to explore the cause and effect relationships between the measured variables through latent values. PLS-PM is effective in both exploring unknown relationships as well as combining large scale data, such as field, physiological, and morphological data, that otherwise is not well described together (Barberán *et al.*, 2014). In addition to running the PLS-PM, 1000 bootstraps were performed to obtain statistical significance and confidence intervals of the path coefficients and the R^2^ values of each latent variable. The path coefficients are the standardized partial regression coefficients (Barberán *et al.*, 2014), and represent the direction and strength of causal relationships of direct effects. Indirect effects are the multiplied coefficients between the predictor variable and the response variable of all possible paths other than the direct effect (Barberán *et al.*, 2014). To determine the best path model, the latent variables were combined using our best understanding of biological relationships, and a general model using all data was generated. The paths between latent variables (LV) were altered until a best-fit model was found. PLS-Predict was then used on the dataset to ensure that the model did not over or under fit the data, and for predictive performance of each manifest variable (MV). This structural model, and not the fit values, was retained for use in predictive modeling of a separate dataset.

PLS-Predict, with the structural model developed as above, was used on a separate dataset to determine the efficacy of the model. Two commercial cultivars, M82 and Lukullus, were used and only the leaf shape values were entered as exogenous variables. The predicted values for each output variable (yield, BRIX, and vegetative mass) were compared to the actual measured values to determine how well the model predicted these variables.

### DNA library construction and sequencing

DNA was extracted using GeneJET Plant Genomic DNA Purification Mini Kit (Thermo Scientific, Waltham, MA, USA) from plants grown for a month and DNA-Seq libraries were prepared based on BrAD-seq (Townsley *et al.*, 2015) with the following modifications: After DNA fragmentation with Covaris E220 (Covaris, Inc. Woburn, MA, USA), the fragmented DNA was end-repaired, A-tailed, and adapter ligated with Y-adapter. Enrichment PCR was then performed with the adapter ligated product as described Townsley *et al.*, 2015. After final library cleanup with AMPure beads (Beckman Coulter, Brea, CA, USA), DNA-Seq libraries were sequenced at Novogene (Novogene, Inc. Sacramento, CA, USA) using the HiSeq 2500 platform (Illumina Inc. San Diego, CA, USA). All data is deposited in SRA (accession: prjna527863).

### Phylogenetic Analyses

To perform phylogenetic analysis, all variants detected by CLC Genomics Workbench 11.0 (CLC Bio, a QIAGEN Company, Aarhus, Denmark) were exported as vcf files. The SNPRelate package for R (Zheng et. al., 2012) was used to determine the variant positions that overlapped between cultivars and then all sequences combined into a single gds file. This file was run through SNPhylo (Lee et. al., 2014) with the following parameters: The linkage disequilibrium was set to 1.0, as we wanted to exclude as few variants as possible based on this factor, minor allele frequency was set to 0.05, and the Missing rate was set to 0.1. One thousand bootstraps were performed for confidence intervals and significance. *S. pimpinellifolium* was used as the outgroup. The bootstrapped output tree was displayed in Mega7 (Kumar, Stecher, and Tamura, 2015).

## Results

### Fruit and Vegetative Traits

To determine the differences in fruit quality between each of the eighteen cultivars, the yield, and fruit BRIX (soluble solids content) were measured over fourteen weeks of the growing season (Fig. S1). For most cultivars the yield remained at or below 5kg of fruit per plant at the time of harvest (Fig. S1a). The exceptions to this were Bloody Butcher, Glacier, Brandywine, Prudens Purple, and Stupice. All had a yield higher than 5kg per plant, with Stupice having the highest yield at approximately 20kg per plant by final harvest (week 23; Fig. S1a). Fruit BRIX remained nearly constant for all cultivars across the growing season (Fig. S1b), with the exception of ABC Potato Leaf, which had a large increase at time of harvest.

To better quantify fruit quality, BRIX and yield were multiplied to obtain a BY value (Grandillo *et al.*, 1999) for each measurement (Fig. 1). The BY value shows a similar trend to yield but identifies cultivars which sacrifice BRIX for yield. Bloody Butcher, Glacier, and Stupice all had a BY value greater than 60, with Stupice reaching near 100 at terminal harvest (Week 23; Fig. 1). The average BY value for harvest weeks (17 to 23) was 16.39, while Bloody Butcher, Brandywine, Glacier, and Prudens Purple had an average at approximately 23 (Table 1). Stupice showed the highest deviation with a mean BY value of 37.86 (Table 1), setting it apart as the highest fruit quality cultivar tested in this study. Stupice maintained a stable BRIX content in its fruit despite the large increase in yield (Fig. S1) which resulted in the large increase in overall fruit quality compared to other cultivars. Vegetative traits such as total biomass and leaf area were measured for the growing season as well (Fig. 2). Figure 2a shows the vegetative biomass and leaf area over the course of the growing season, which remain stably linked indicating that overall leaf area increase contributed to increased biomass of the plant. This trend appears common in heirloom tomatoes but is different in commercial tomatoes, which have determinate growth (Pnueli, *et al.*, 1998).

**Table 1.** Mean BY values for the eighteen heirloom cultivars with standard error, number of plants sampled, and the final row containing the mean of all lines measured during the harvest time period.

| Genotype | Mean BY | Std Error | N |
| --- | --- | --- | --- |
| ABC Potato Leaf | 17.99 | 5.47 | 12.00 |
| Aunt Ginny's Purple | 2.34 | 0.68 | 10.00 |
| Bloody Butcher | 21.78 | 8.08 | 11.00 |
| Blue Ridge Mountain | 10.79 | 3.33 | 5.00 |
| Brandywine | 23.40 | 10.42 | 12.00 |
| Burwood's Prize | 15.79 | 7.62 | 7.00 |
| Depp's Pink Firefly | 12.35 | 7.79 | 3.00 |
| Glacier | 24.39 | 11.42 | 6.00 |
| Green Pearl | 12.51 | 11.35 | 3.00 |
| Jerry's German Pink | 16.62 | 4.07 | 8.00 |
| Marianna's Peace | 6.01 | 2.88 | 6.00 |
| Matina | 9.78 | 5.04 | 6.00 |
| Prudens Purple | 24.59 | 5.83 | 11.00 |
| Rockingham | 8.53 | 3.31 | 9.00 |
| Schelicauski | 10.39 | 2.35 | 7.00 |
| Soldacki | 6.82 | 2.22 | 7.00 |
| Stupice | 37.86 | 16.93 | 9.00 |
| Valena Pink | 14.84 | 5.09 | 11.00 |
| Week 17 to 23 Mean | 16.39 | 1.91 | 143.00 |

**Fig. 1.**
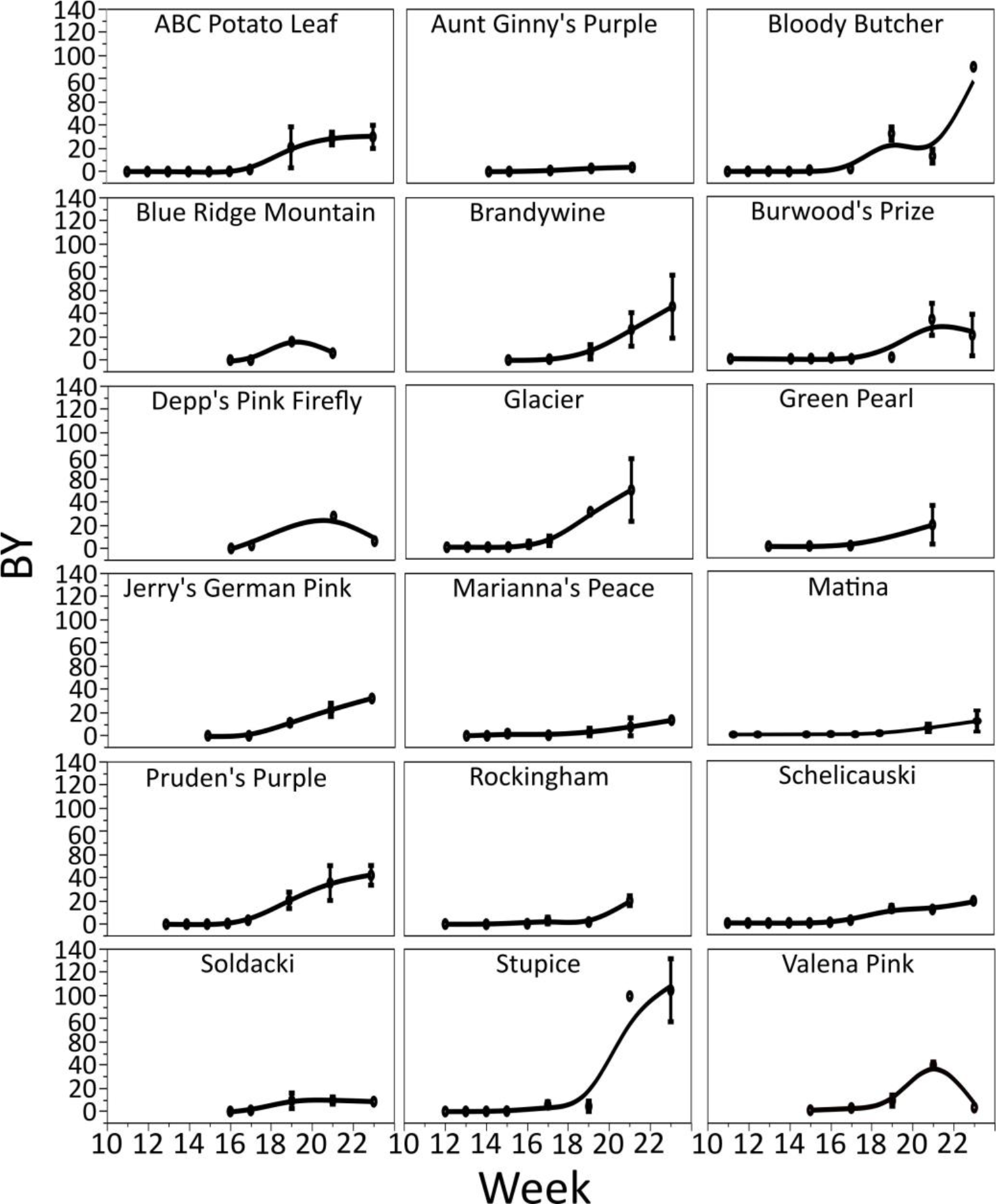
BRIX by Yield Index of eighteen heirloom cultivars. Potato Leaf Morph heirloom tomato cultivars were grown in the field and the fruit BRIX and yield measured over a fourteen-week growing period. The fruit BRIX and Yield were then multiplied together to obtain the BY value, giving a better indicator of overall fruit quality. Stand out cultivars were Bloody Butcher, Brandywine, Glacier, Prudens Purple, and Stupice all of which obtained a BY value greater than twenty. The average BY value during harvest weeks (week 17 to 23) was 16.39. The values shown are the mean BY, and error bars represent standard error.

**Fig. 2.**
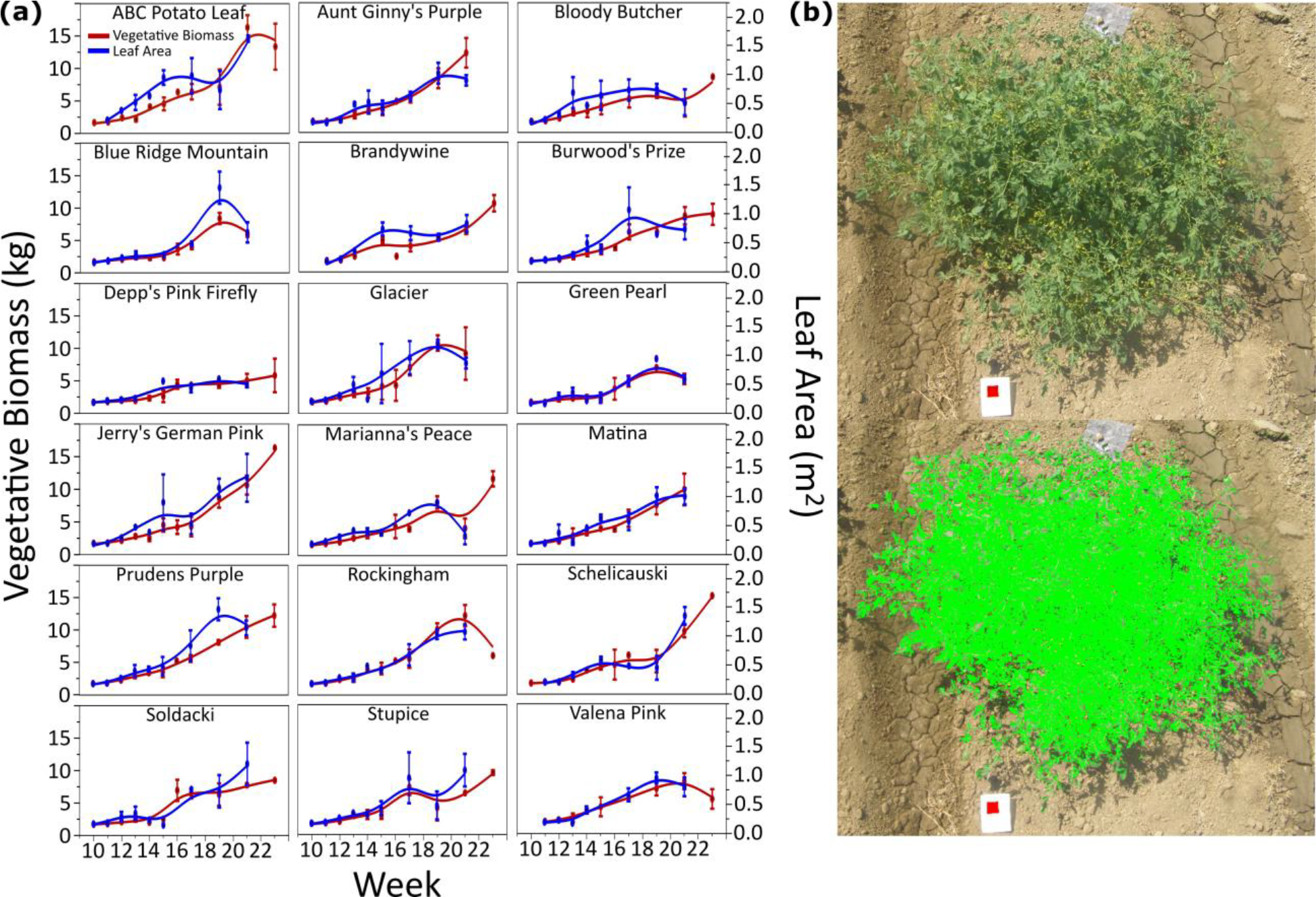
Vegetative biomass and leaf area of heirloom tomatoes. Over the fourteen-week growing period the total vegetative biomass (fresh weight) and leaf area were measured. (a) The mean measurements for both total biomass and leaf area for each cultivar. Error bars represent standard error. Leaf area mostly followed vegetative biomass, increasing incrementally with biomass, however in Burwood’s Prize the leaf area levels out around week 17 and does not increase with increase biomass. (b) Represents the method with which leaf area was obtained. Overhead photos were taken of each plant, with a red square of known size, and then using (software) the green area exposed to sunlight was calculated.

### Photosynthesis

As photosynthesis is the primary source of sugar production in plants, a time course for photosynthesis, stomatal conductance (*gst*), intercepted photosynthetically active radiation (PARi), and ɸPS2, was performed on all cultivars using a LI-6400XT (Fig. 3; LI-COR, Lincoln, NE). Figure 3a shows photosynthesis by *gst* and the trend is similar among all cultivars with photosynthesis reaching a maximum rate after 0.6 to 0.8 *gst*, which is a standard response curve (Gilbert *et al.*, 2011). Valena Pink, Rockingham, and Marianna’s Peace however show a more linear relationship suggesting that given a higher *gst* the photosynthetic rate would continue to increase. Figure 3b and 3c show the PARi and ɸPS2 respectively. All cultivars reached a PARi of greater than 1200 over the time course, though when this peak was reached varied by cultivar, due mostly to the duration of growth specific to each cultivar. ɸPS2 had an overall downward trend across the season, as the amount of light used in photo system II decreased with age. This corresponds well to the increase in vegetative biomass (Fig 2a), and the increased PARi (Fig. 3). Individual leaf contribution to overall photosynthesis, and therefore, photons used in PSII decreases as the leaf area of the plant increases.

**Fig. 3.**
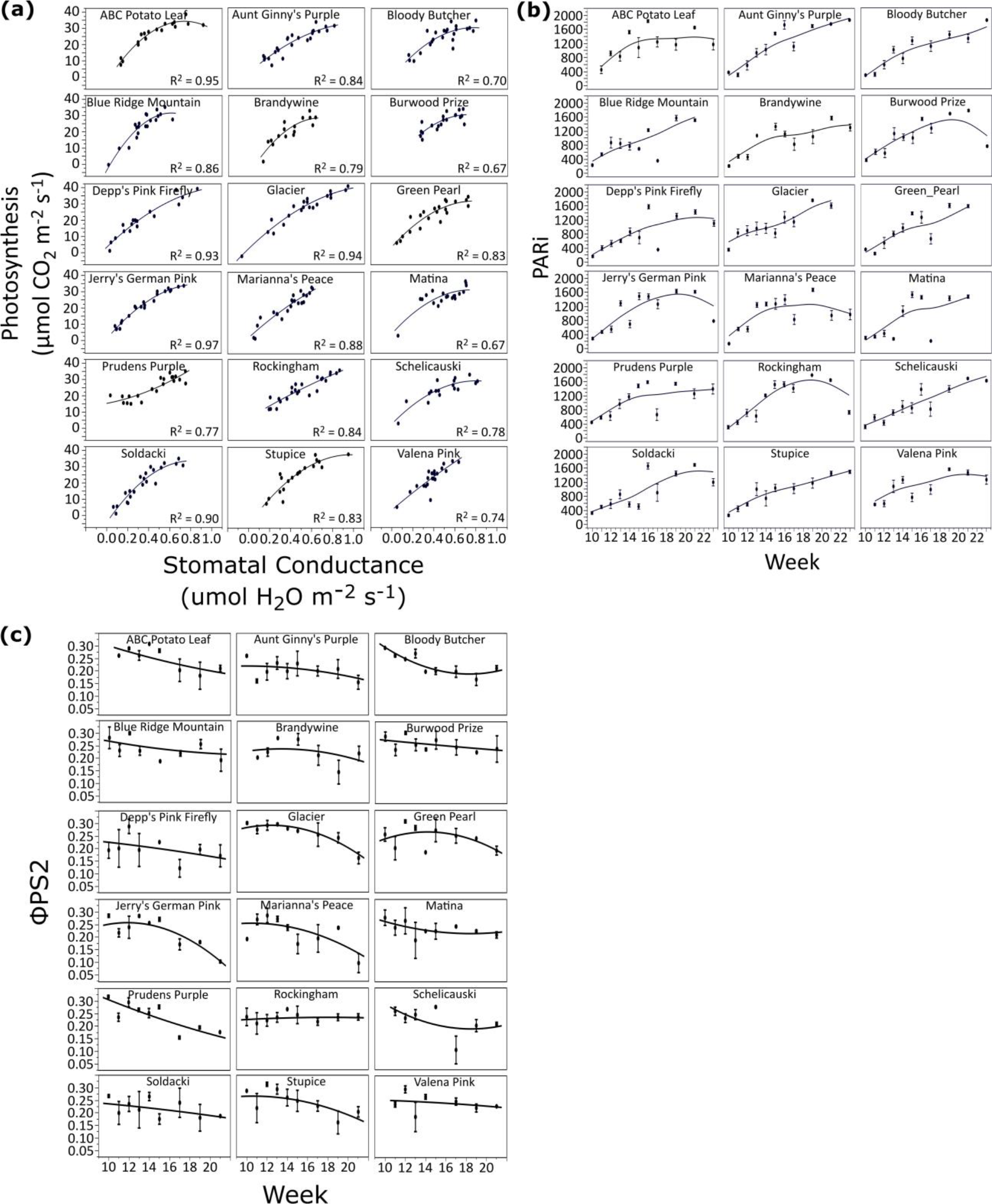
LICOR LI-6400XT and LQA 2857 measurements. (a) Photosynthetic rate (*A*) and stomatal conductance (*gst*) were measured using the LI-6400XT for two leaves per plant (bottom and top of plant). The correlation between *A* and *gst* is shown, with the majority of lines having a logarithmic curve. Rockingham and Valena Pink have a more linear relationship, and Prudens Purple shows a near exponential increase. (b) PARi was measured using the LQA-2857 and (quantum sensor here) by placing the line sensor beneath the plant and the quantum sensor above and taking four measurements per plant and averaging the result. All plants show a steady increase in PARi over time corresponding to increased vegetative biomass and leaf area. (ɸPSII was measured with the LI-6400XT (fluorescence head info). Values are the mean measurements over time and error bars represent standard error. The overall ɸPSII decreases overtime, with the only exception being Rockingham which remains stable over the entire growing season.

Because of this trend we calculated the whole plant photosynthesis and *gst* to capture the total rates and not just specifically measured leaves (Fig. S2). The trend is linear for photosynthesis versus *gst* when the whole plant exposed green area is incorporated, compared to Figure 3a where the trend is more logarithmic. Rockingham, the worst performing cultivar, shows a logarithmic relationship at the whole plant level as compared to the linear relationship at the individual leaf level.

### Leaf Shape and *C*-Locus Mutations

Leaf shape was shown to be strongly correlated with fruit BRIX and sugar accumulation in an introgression population, with rounder and more circular leaves having the highest sugar content in their fruit (Chitwood *et. al.*, 2013). How leaf shape contributed to fruit BRIX was unclear, as shape and size of leaves do directly impact photosynthesis (Sarlikioti *et al.*, 2011), but direct links between leaf shape and fruit quality appear lacking. Our heirloom cultivars displayed a wide array of leaf shapes, from very large and narrow to small and rounder. To understand if this range of leaf shapes had any impact on the overall fruit quality we measured leaflet shape and size for approximately 3733 leaflets using the MOMOCS R package and Fiji (Bonhomme *et al.*, 2014; Schindelin *et al.*, 2012). Figure 4 shows the resultant PCs of all primary leaflets (terminal and lateral) measured and their relationship to traditional shape measures. PC1 contributes 78% of all variation found in the population and is tightly correlated with leaflet size (R^2^=0.99), indicating that size was the largest source of variation among the heirloom leaflets (Fig. 4). PC2 and PC4, while having no traditional shape measure correlation, indicate the left and right-handedness of the lateral primary leaflets, as these leaflets are mirror images of each other and therefore this measure describes the overall variation in leaf symmetry (Fig. 4a; Chitwood *et al.*, 2013). PC3 accounts for 3.8% of all variation, but has a strong correlation with aspect ratio, or the width divided by the length of the leaflet, with an R^2^ of 0.8 (Fig. 4). PC3 therefore represents the roundness or narrowness of the leaflets, one aspect previously shown to be linked to fruit quality (Chitwood *et. al.*, 2013). PC5 accounts for only 1.6% of all variation and represents the base to tip ratio of the leaflet (Fig. 4a). This variation is interesting as it has no effect on aspect ratio (width and length remain the same), but still represents a change in leaflet shape.

**Fig. 4.**
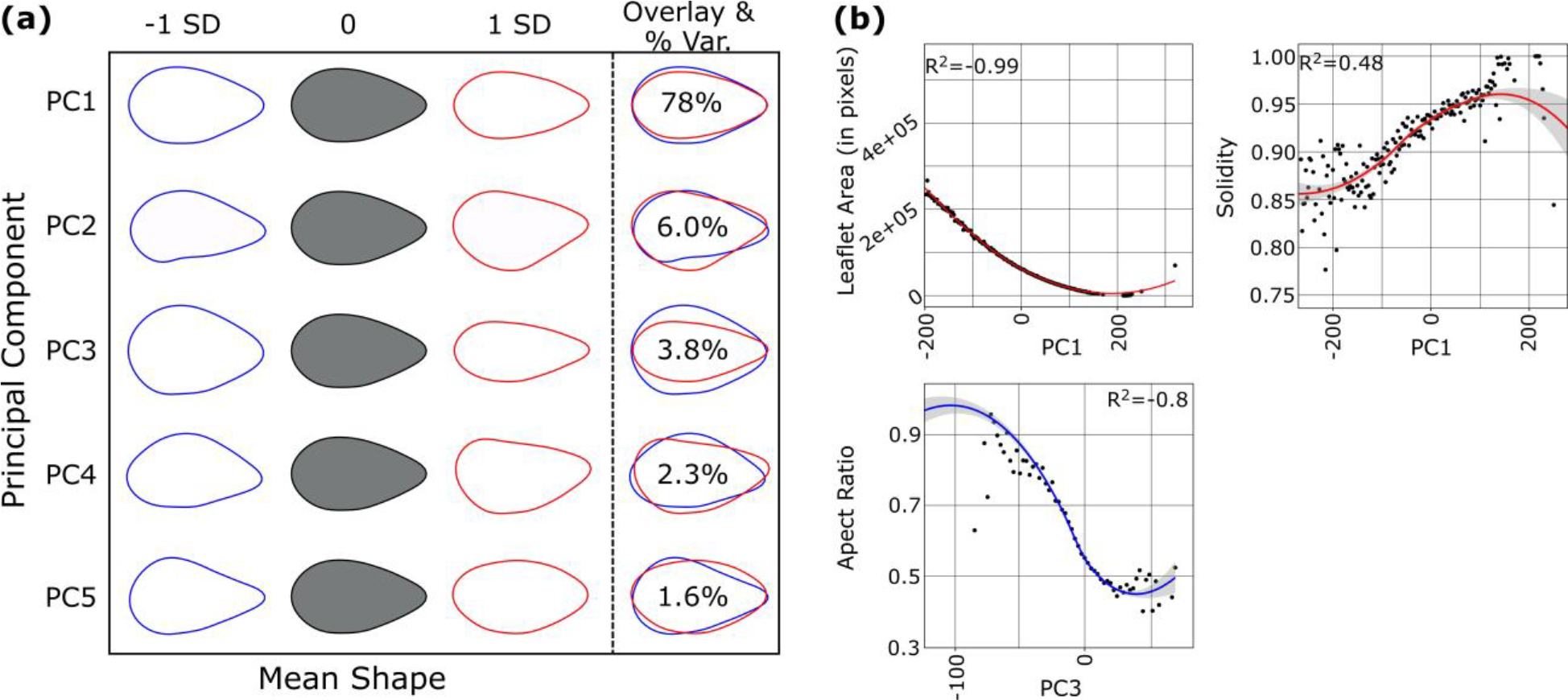
Leaf shape analysis of heirloom tomatoes. (a) The calculated PC values for 3733 leaflets are shown. PC1, PC3, and PC5 account for 83.4% of all variance found in the leaves, with PC1 having 78% of the variance. (b) Correlations of PC values to traditional measurements. PC1 correlates with leaflet size (R^2^=0.99), and solidity (R^2^=0.48) which is an indicator of how serrated or lobed a leaflet is. Higher values of solidity indicate a smoother less lobed leaflet. Size is represented here in total number of pixels for correlation purposes. PC5 does not have a traditional shape measure associated with it, but does represent the tip to base ratio of the leaflet. PC2 and PC4 are inversions of the lateral primary leaflets which are mirror images of each other represented by these PCs.

The heirloom cultivars analyzed all have broader, smoother leaves and typically lack the serration and lobes seen in other tomato varieties (Goldman, 2008; http://tatianastomatobase.com/wiki/Main_Page). However, despite this they had a wide range of leaf shape and size as illustrated in the leaf shape analysis (Fig. 4). The classical *potato leaf* mutation (abbreviated to *c*) is caused by a 5kb transposable element (RIDER; Jiang *et al.*, 2009; Jiang *et al.*, 2012) inserted into the third exon of the *C* locus (Solyc06g074910; Busch *et al.*, 2011). To determine if this locus harbored mutations in the selected lines, a subset of the higher performing cultivars were selected for whole genome sequencing (WGS) analysis. Other mutations at the *C* locus have been described, and cause varied leaf shape, (Busch *et al.*, 2011). We reasoned that the selected heirlooms may have different mutations in the *C*-locus, which, along with their diverse breeding histories, could account for the varied leaf shapes seen in these lines. Figure 5a shows the location of the mutations found in the *C* Locus in these select lines. While the full Rider insertion could not be directly determined as the reference genome lacks this insertion, overhangs on reads in the third exon matched the Rider TE sequence (Fig. 5a, S4). It is possible that different sizes and fragments of Rider are present in different cultivars, as the length and sequence of the overhangs varied (Fig. S4). The identified Rider sequences were present in all but two of the sequenced lines, with both Prudens Purple and Glacier missing this element. No mutations were found in Glacier despite it having a rounder leaflets, though these were smaller in size with higher overall leaf complexity (Fig. 4, 5, S3). Prudens Purple did not have the RIDER insertion but did have a novel single base pair substitution in the first exon outside the MYB/SANT conserved domain which results in the amino acid change P42R (Fig. 5a). We analyzed this mutation using the Protein Variance Analyzer (PROVEAN; Choi, 2012; Choi *et. al.*, 2012; Choi and Chan, 2015), and found that it is predicted to be deleterious to the protein with a value of −8.454 (threshold set at −2.5), predicted to result in either a non-functional or partially functional protein (Choi, 2012; Choi *et. al.*, 2012; Choi and Chan, 2015). Based on leaf shape analysis, Prudens Purple shows a Potato Leaf like phenotype (Fig. S5), although it differs slightly from the classical Potato Leaf shape seen in the reference allele and is reminiscent of the other mutations in *C* that have varying leaf shapes (Busch *et al.*, 2011). These data demonstrate that different mutations in *C* give rise to a range of leaf shapes, resulting in the wide leaf variation seen among these cultivars.

**Fig. 5.**
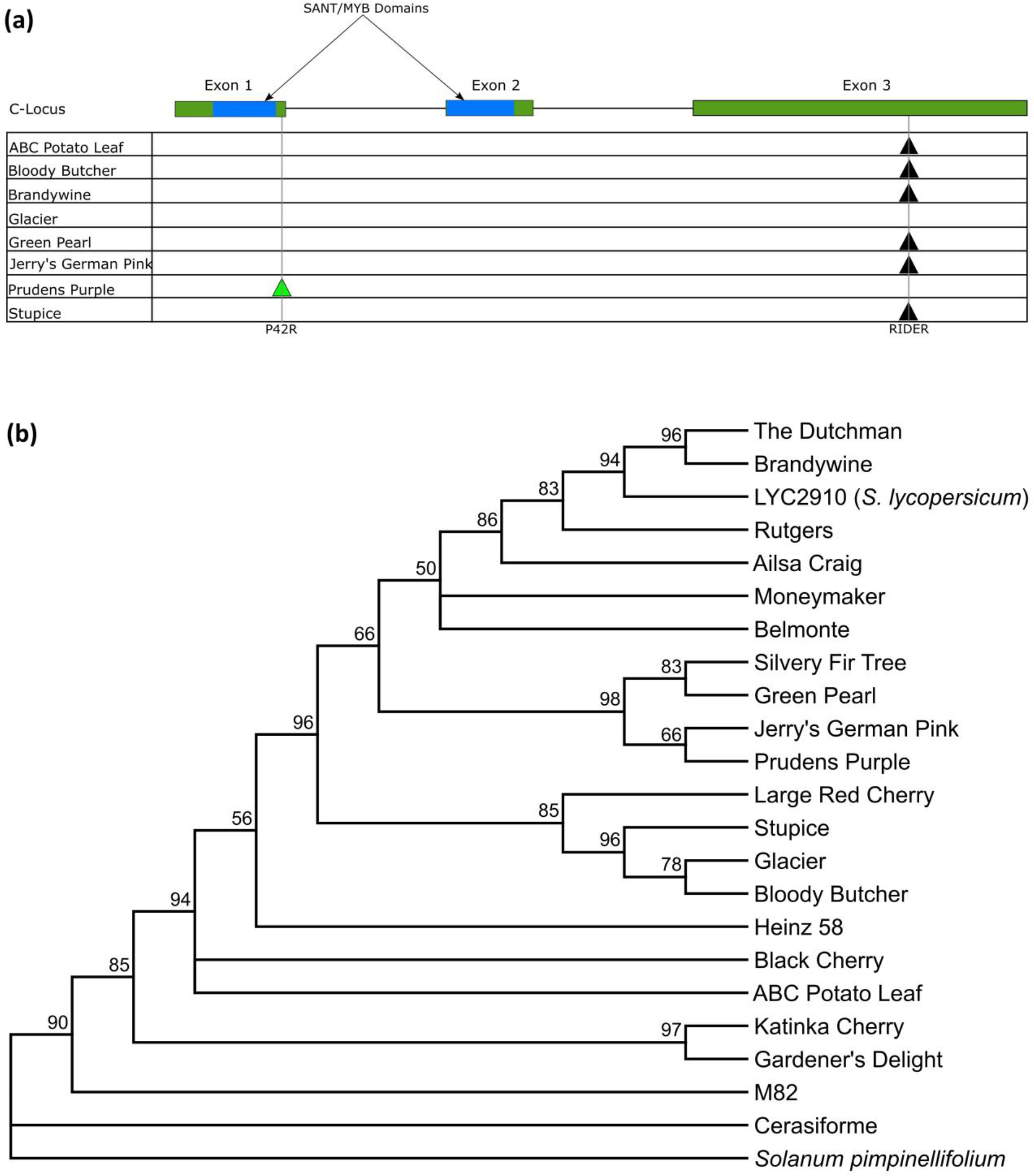
*C*-Locus mapping and WGS based Phylogeny. (a) Eight heirloom cultivars were sequenced using WGS and the mutations found in the *C*-Locus are shown here. Black arrows indicate the position of the RIDER TE insert, and the green arrow indicates the single base pair mutation and its resultant amino acid swap. Glacier had no mutations found in the *C*-Locus. (b) Using variants from the WGS sequencing a phylogeny was generated for the eight heirloom cultivars and fourteen additional lines. M82 was sequenced by WGS by us, while the remaining sequences were obtained from the 150 Genomes Project (Aflitos *et al.*, 2014).

### Phylogeny

We had little or no information on the breeding history of many of the heirlooms used in this study. Pedigrees would likely inform the overall leaf shape in addition to the source of the *C-*Locus mutations. To elucidate relationships among these cultivars we used the WGS data from the select cultivars as well as WGS data obtained from the 150 Genomes Project (Aflitos *et al.*, 2014) to assemble a phylogeny (Fig. 5b). The phylogeny includes several commercial cultivars, heirloom commercial cultivars, *Solanum pimpinellifolium*, and *Solanum lycopersicum var. cerasiforme*. ABC Potato Leaf is unusual in that it appears to be more closely related to the wild relatives than to other Potato Leaf heirloom cultivars analyzed here (Fig. 5b). The non-Potato Leaf heirlooms form a clade separate from the Potato Leaf Heirlooms with the exception of Brandywine, which is most closely related to other commercial cultivars. Stupice, Glacier, and Bloody Butcher are closely related in this phylogeny. This corresponds to their often being listed as closely related (Goldman, 2008) and is congruent with phenotypic similarities in fruit size and leaf shape. Glacier as noted above does not contain a mutation in the *C*-Locus and has a much higher leaf complexity. However, it has a rounder leaf shape than most other cultivars observed, (Fig. 4, 5a, S3). Bloody Butcher and Stupice both have the RIDER insertion in the 3^rd^ exon at the *C* locus (Fig. 5a), suggesting presence of other modifiers to leaf shape which may have been selected for during the breeding of Glacier. A similar situation is seen in Prudens Purple (Fig. 5b), which is most closely related to Green Pearl. These two heirlooms carry distinct mutations at the *C* locus. Green Pearl has the RIDER TE in the 3^rd^ exon (Fig. 5a) and a classical Potato Leaf Morph, whereas Prudens Purple has a novel single base pair substitution in the first exon leading to a deleterious effect on protein function. While these cultivars come from a similar region of Eastern Europe (Fig. 5b; Goldman, 2008), our data suggest they have a complex breeding history. Lack of clear breeding history for the potato leaf phenotype within the phylogeny, suggests multiple attempts to breed this trait into cultivars. Potato leaf cultivars have been suggested to increase disease resistance compared to regular leaf varieties (Male, 1999) and may have been selected for this reason. Improved fruit quality in potato leaf morphs may have been attributed to increased resistance more than the leaf shape itself.

### Partial Least-Squares Path Modeling

When doing large-scale field studies, it is difficult to understand how all the collected data points relate to each other, and what the causative relationships are (Granier and Vile, 2014). To decipher how all the physiological and morphological traits measured related to each other, we performed Partial Least-Squares Path Modeling (PLS-PM) using SmartPLS3.0 (Ringle *et al.*, 2015), which gives weighted causative paths with bootstrapping for confidence and significance values. In PLS-PM, each LV (such as Leaf Shape) is a composite value of its associated Manifest variables (determined through correlations) and forms an outer model (Table S1). The inner model consists of the connections between LVs, with R^2^ values indicating the degree to which each endogenous LV is described by the connections to it (Table S2). Here, the only exogenous LV is Leaf Shape, as it has only its associated MVs and is descriptive of other LVs. Some LVs (Photosynthesis) are described by other LVs within the model (such as gas exchange, light input in the case of photosynthesis). When the value of a causative LV (such as leaf shape) increases the corresponding connected LV/s change in accordance with their relationship with the causative variable. Similarly, in the outer model, changes in MVs reflect a change in their LV, and thus connect the outer model with MVs (measured values) to the inner model of LVs. For instance, the MV PC3 has a negative correlation with the LV Leaf Shape (Table S1, Fig. S6), so as the value of PC3 decreases, it reflects as a corresponding increase in LV Leaf Shape (Fig. S6). This change is represented as an increase in the roundness of the leaf. This then corresponds to a positive change in Yield (LV), which is in turn a reflection of Fruit Biomass (MV) (Fig. S6).

The model indicates that photosynthesis has a strong positive influence on both fruit BRIX and vegetative biomass but has a negative impact on fruit yield. As photosynthetic rates increase (along with light capture and gas exchange) fruit BRIX increases, but at the sacrifice of yield, an inverse relationship which has long been known (Fig. 6; Eshed and Zamir, 1994; Zanor *et al.*, 2009; Chitwood *et al.*, 2013; Fridman *et al.*, 2000; Osorio *et al.*, 2014; Lytovchenko *et al.*, 2011). Leaf shape has a negative relationship with vegetative biomass, which corresponds to the decreased leaf complexity with the Potato Leaf Morph (Fig. 5, 6, S3). However, leaf shape has a strong positive influence on both fruit BRIX and yield (Fig. 6), suggesting that leaf shape influences fruit quality as seen previously by Chitwood and coworkers (2013). The effect of leaf shape on fruit quality does not work through leaf sugar, as this correlation was not significant.

**Fig. 6.**
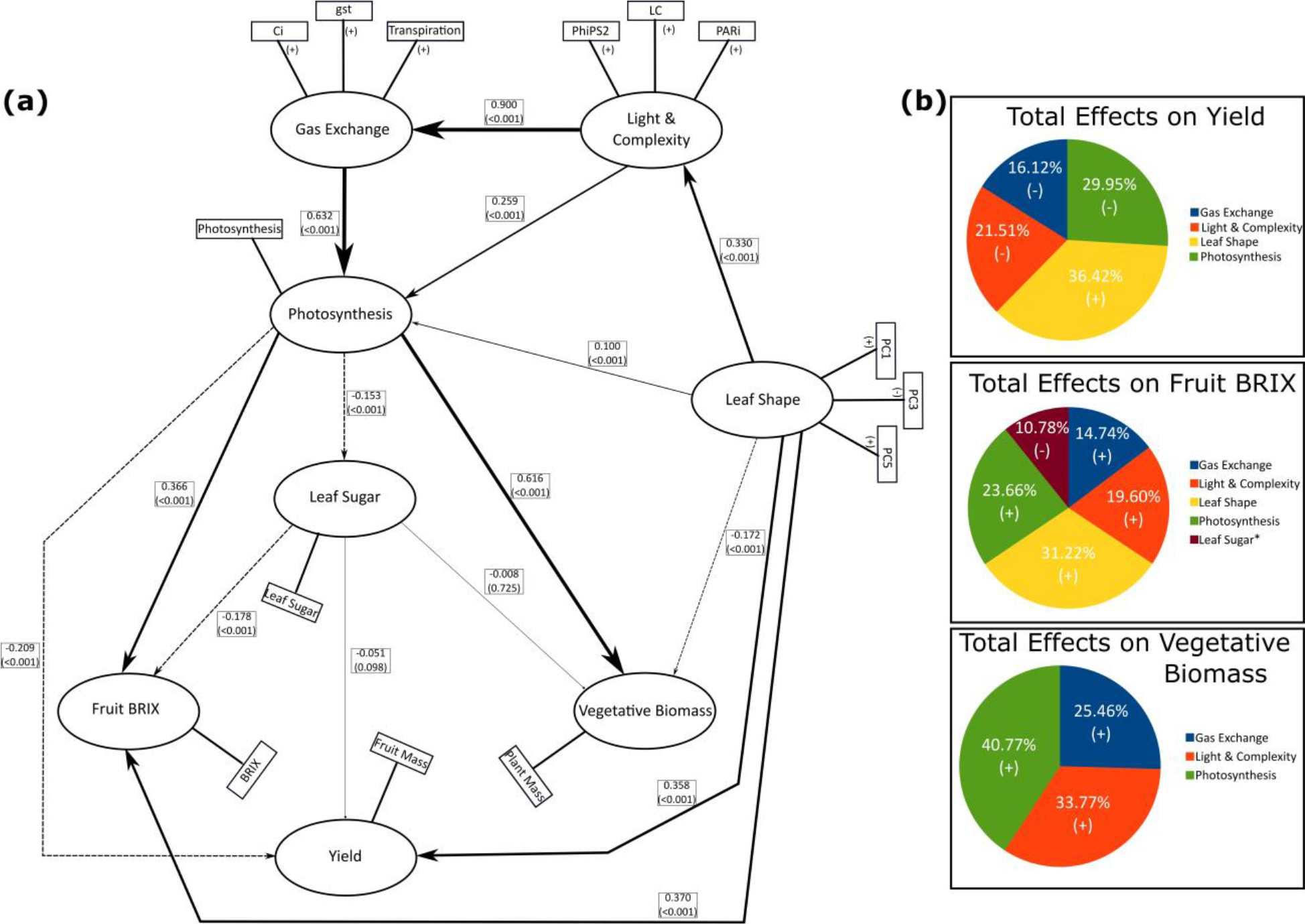
Partial Least Squares-Path Modeling (PLS-PM) of all collected physiological and morphological data. (a) The finalized version of the PLS-PM using SmartPLS3.0 is shown. Traits within the circles are Latent Variables (LV’s) which represent the measurements in rectangles, the Manifest Variables (MV’s). The (+) and (-) next to each MV represents its relationship to its LV. PC1 and PC5 are positively correlated to the LV Leaf Shape, while PC3 is negatively correlated. This corresponds to a rounder leaflet shape (-PC3, +PC5). Solid lines indicate a positive correlation and dashed lines indicate a negative correlation between LV’s. The size of the arrow indicates path weight, which is listed next to the path along with the p-value. Blunted lines indicate non-significant connections. Photosynthesis has a strong positive correlation with Fruit BRIX and Vegetative Biomass but has a negative correlation with Yield. Leaf Sugar content has no significant impact on Vegetative Biomass or Yield, but is negatively correlated with Fruit BRIX, indicating that a lower Leaf Sugar content corresponds to an increased Fruit BRIX. Leaf Shape has a positive correlation with both Yield and Fruit BRIX, indicating that Leaf Shape may play a role in distribution of photosynthate. (b) The total effects of different LV’s on the outputs of the PLS-PM: Yield, Fruit BRIX, and Vegetative Biomass. For both Yield and Fruit BRIX Leaf Shape has the strongest effect, followed by photosynthesis. (+) indicates a positive effect while (-) indicates a negative effect. Percentages are the proportion of path weights contributing to each output.

Figure 6b displays the effect of each trait on the overall output of the plants (fruit BRIX, yield, and vegetative biomass). Leaf shape has no strong contribution to vegetative biomass. Although shape shows a negative relationship with biomass, this influence is minimal when compared to photosynthesis (Fig. 6b). However, leaf shape shows the largest influence on both yield and fruit BRIX with photosynthesis second, and is the only positive contributor to yield (Fig. 6b). This positive correlation is from rounder, Potato Leaf Morph-like leaves while narrower leaves have the opposite effect (Fig. 6a) based on the PC contributions to leaf shape. The negative effect of photosynthesis on tomato fruit yield and the strong contribution of leaf shape to yield and BRIX are novel findings which run counter to the interpretation of fruit quality improvement, as increased photoassimilate should result in more available sucrose to stronger sinks such as fruit (Osorio *et al.*, 2014).

To test the model performance we used PLS-Predict on the entire heirloom dataset used to build the structural model. Table S3 shows the MAPE (Mean Absolute Percent Error) and Q2 value for the complete model. We also used part of the dataset that included ABC Potato Leaf, and Aunt Ginny’s Purple in a similar analysis (Table S3). The complete model has approximately 20% to 30% error for each LV, which is expected given the diversity of genotypes in the dataset, with fruit weight giving the highest MAPE at 93.2% (Table S3). The Q2 value for most variables is positive and shows that they have relevance in the predictive performance, with the exception of Leaf Sugar, which is slightly negative (Table S3). In the case of ABC Potato Leaf and Aunt Ginny’s Purple, two lines selected randomly to test the model on individual cultivars, a significant increase in Q2 and decrease in MAPE is seen for all LVs except Leaf Sugar (Table S3). This indicates that the model is substantially stronger in predictive performance for individual cultivars, but also predicts well with the complete model.

To evaluate the predictive performance of our model on additional datasets we used data from two other cultivars grown in the same field, M82 and Lukullus, that were not used to construct the model. PLS-Predict was used in SmartPLS 3.0, along with the structural model constructed using the heirloom cultivars, to test the model performance by use of training sets and hold out samples, both taken from the M82/Lukullus dataset. By using the Leaf Shape PC values we were able to compare the predicted mean values for the remaining manifest variables (MV), or the predicted measured values, against the actual measured values and evaluate the relative performance of the model. Tables 2 and 3 show the results for M82 and Lukullus respectively. PC values for leaf shape are not included as they are input variables and used for predicting the other values. For M82 the predicted median values compared to the actual median values showed under 1% difference for all except for leaf complexity, which had a percent difference of −8.42% (Table 2). This indicates that the model was under predicting the leaf complexity of M82 by ~8 percentage points. Lukullus predicted values were also under 1% difference except for leaf complexity and stomatal conductance which varied by −2.56% and 1.31% respectively (Table 3). In addition to the predicted values PLS-Predict also tests the model performance and reports the Root Mean Square Error (RMSE), Mean Absolute Error (MAE), and Mean Absolute Percent Error (MAPE) for each of the MVs tested (Table 2, 3). The MAPE shows the accuracy of the predictions with lower percentages representing better performance. Leaf complexity for both cultivars showed the largest MAPE of 201.2% and 26.5% in M82 and Lukullus respectively (Table 2, 3). The M82 MAPE indicates the model does not predict leaf complexity well for mid-level complexities such as 18 but does improve at high end leaf complexities near 40 (Table 2, 3). Most heirloom cultivars had low leaf complexities (Fig. S3) potentially explaining the poor performance in predicting leaf complexity for M82. We found that leaf complexity does not impact yield or BRIX, and only impacts vegetative biomass, so this inaccuracy would only impact vegetative output predictions by the model. Lukullus has indeterminate growth like the heirlooms analyzed here, but M82 is determinate, however the predictive accuracy of the model was still good, indicating its usefulness in assessing field performance of other tomato cultivars.

**Table 2.** Predicted and actual median values for M82 of MV’s in the PLS-Path Model, with error rates for model accuracy.

| MV | Predicted Median | Actual Median | Percent Difference | RMSE | MAE | MAPE |
| --- | --- | --- | --- | --- | --- | --- |
| Stomatal Conductance | 0.152 | 0.152 | 0.14% | 0.014 | 0.010 | 6.92 |
| Internal CO2 | 72.098 | 71.623 | 0.66% | 7.038 | 5.897 | 8.21 |
| Transpiration (mmols) | 2.720 | 2.736 | -0.60% | 0.231 | 0.189 | 7.07 |
| PhiPS2 | 0.071 | 0.070 | 0.96% | 0.006 | 0.004 | 6.08 |
| PARi | 952.922 | 956.761 | -0.40% | 15.947 | 12.462 | 1.31 |
| Photosynthesis | 7.575 | 7.531 | 0.59% | 0.607 | 0.457 | 6.12 |
| Leaf Sugar | 2.723 | 2.725 | -0.08% | 0.092 | 0.074 | 2.75 |
| Complexity | 16.310 | 17.809 | -8.42% | 9.719 | 7.806 | 201.28 |
| Plant Weight | 2.864 | 2.888 | -0.82% | 0.427 | 0.339 | 12.25 |
| Fruit Weight | 1.246 | 1.236 | 0.76% | 0.209 | 0.163 | 13.21 |
| BRIX | 4.134 | 4.136 | -0.04% | 0.038 | 0.032 | 0.76 |

RMSE, Root Mean Square Error; MAE, Mean Absolute Error; MAPE, Mean Absolute Percent Error.

**Table 3.**
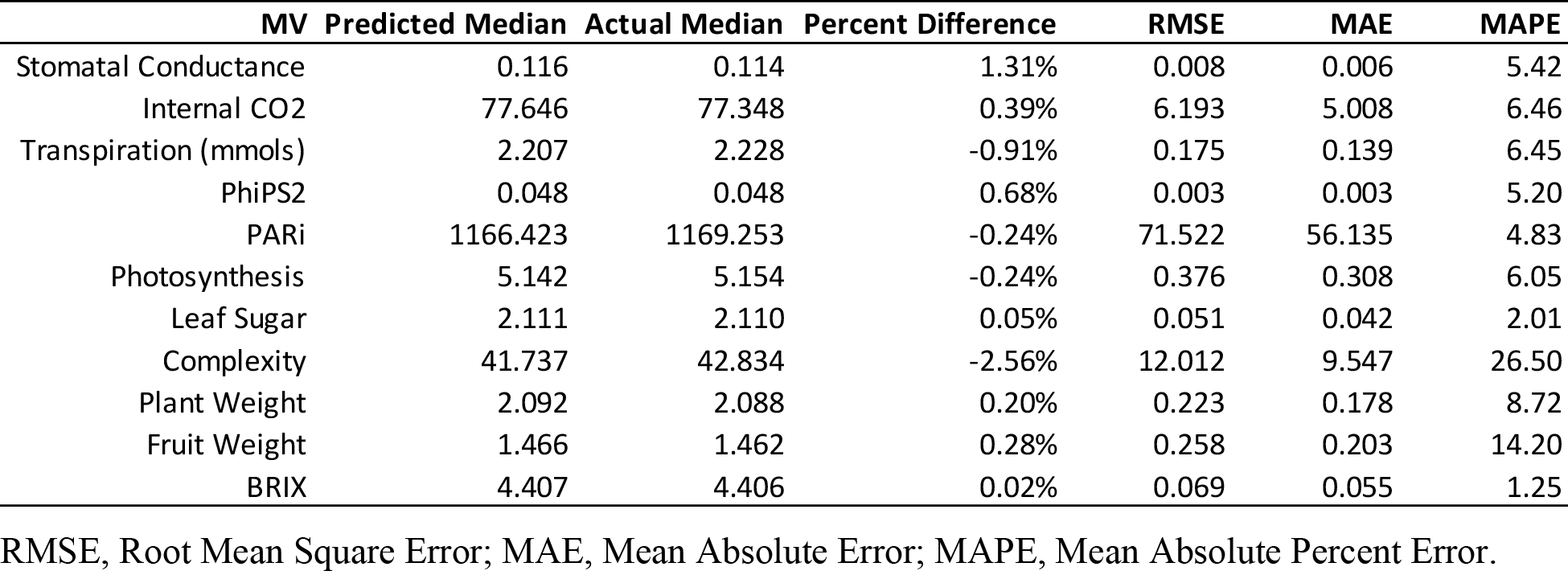
Predicted and actual median values for Lukullus of MV’s in the PLS-Path Model, with error rates for model performance evaluation.

| MV | Predicted Median | Actual Median | Percent Difference | RMSE | MAE | MAPE |
| --- | --- | --- | --- | --- | --- | --- |
| Stomatal Conductance | 0.116 | 0.114 | 1.31% | 0.008 | 0.006 | 5.42 |
| Internal CO2 | 77.646 | 77.348 | 0.39% | 6.193 | 5.008 | 6.46 |
| Transpiration (mmols) | 2.207 | 2.228 | -0.91% | 0.175 | 0.139 | 6.45 |
| PhiPS2 | 0.048 | 0.048 | 0.68% | 0.003 | 0.003 | 5.20 |
| PARI | 1166.423 | 1169.253 | -0.24% | 71.522 | 56.135 | 4.83 |
| Photosynthesis | 5.142 | 5.154 | -0.24% | 0.376 | 0.308 | 6.05 |
| Leaf Sugar | 2.111 | 2.110 | 0.05% | 0.051 | 0.042 | 2.01 |
| Complexity | 41.737 | 42.834 | -2.56% | 12.012 | 9.547 | 26.50 |
| Plant Weight | 2.092 | 2.088 | 0.20% | 0.223 | 0.178 | 8.72 |
| Fruit Weight | 1.466 | 1.462 | 0.28% | 0.258 | 0.203 | 14.20 |
| BRIX | 4.407 | 4.406 | 0.02% | 0.069 | 0.055 | 1.25 |

RMSE, Root Mean Square Error; MAE, Mean Absolute Error; MAPE, Mean Absolute Percent Error.

## Discussion

The primary focus of crop improvement has been on fruit traits (sink) and photosynthesis (source), with some studies focusing on how sugars are moved from source to sink. Despite heirloom varieties with the Potato Leaf Morph being prized for fruit quality by the gardening community, vegetative traits such as leaf shape have been relatively ignored in breeding efforts. In this study we investigated the role of leaf shape on fruit quality by measuring both input traits such as photosynthesis, leaf shape, leaf complexity, and output traits such as yield, BRIX, and vegetative biomass, for eighteen heirloom cultivars. All these cultivars were classified as Potato Leaf, but varied greatly in their leaf shapes, development, and fruit quality (Fig. 1, 2, 4). We found that these lines don’t vary significantly in overall photosynthetic capacity, or their usage of light when available (Fig. 3), suggesting that the variation in BY (Fig. 1) among these cultivars was not due to improved/decreased photosynthetic capacity. While our measurements for photosynthesis don’t show significant difference when PAR is available, the PARi differed between cultivars based on their growth patterns (Fig. 2b, 3b). All cultivars exceeded 1200 μmols m^−2^ s^−1^ of PARi but varied in the later weeks between 1200 and 2000 μmols m^−2^ s^−1^ (Fig. 3b).

Combining multiple complex physiological and morphological measurements into informative relationships has proven difficult and has limited our understanding of how these different traits impact each other (Granier and Vile, 2014). Focusing on any one part such as photosynthesis, or fruit sink strength, while providing improvements (Zanor *et al.*, 2009), occurs at the expense of a comprehensive understanding of the overall relationships between these traits. Analyzing the individual PCs revealed significant differences in leaf shape among the heirloom cultivars, with several having stronger Potato Leaf Morphs (Fig. 4, S5) and higher BY values (Fig. 1), with some correlation between these traits.

We used Partial Least Squares-Path Modeling to combine all these measured traits, using the modeled final harvest data as input to find causative relationships (Fig. 6a). Strong relationships between gas exchange, light, and photosynthesis were expected, along with a strong positive effect of photosynthesis on vegetative biomass (Fig. 6a, b). Photosynthesis has a strong positive effect on fruit BRIX, both directly and indirectly (Fig. 6a). Increased photosynthesis results in lowered leaf sugar content, and a concomitant increase in fruit BRIX. It is possible that increased sugar production from photosynthesis results in higher rates of transport of sugars out of the leaves and into sinks. The mechanisms that regulate source-sink relations and sugar distribution are still not fully understood on a whole plant physiological level (Osorio *et al.*, 2014), however based on our model, increased photosynthesis negatively impacts total yield (Fig. 6a, b, 7). While photosynthesis does lead to increased sugar production and is shown in our model to drive higher sugar content within existing fruit, it does not provide a means to increase yield. Leaf shape, specifically rounder, less lobed leaves have a positive effect on both fruit BRIX and yield (Fig. 6a, b, 7). Of all the factors measured here, only leaf shape positively influenced yield, with other paths having negative influences (Fig. 6b). Rounder leaves still drive slightly increased photosynthesis indicated by the thin arrow (Fig. 7a), which results in increased fruit BRIX. This path should also result in decreased yield. However, leaf shape has a strong positive and direct correlation with yield that overcomes the negative impact of photosynthesis and leads to increased yield as well as BRIX (Fig. 7a). Conversely, with narrow leaflets there is a small negative impact on photosynthesis which should result in increased yield, but narrow leaves have a direct negative impact on yield which is stronger than the photosynthetic pathway (Fig. 7b). The strong causative relationship between leaf shape, fruit BRIX, and yield, suggests that leaf shape impacts both high fruit BRIX as well as increased number of fruits, likely by modulating sugar distribution, therefore bypassing the direct impacts of photosynthesis itself (Fig. 7). How leaf shape affects this distribution is unclear, as it does not act directly through leaf sugar, or through strong regulation of photosynthesis to improve yield (Fig. 7).

**Fig. 7.**
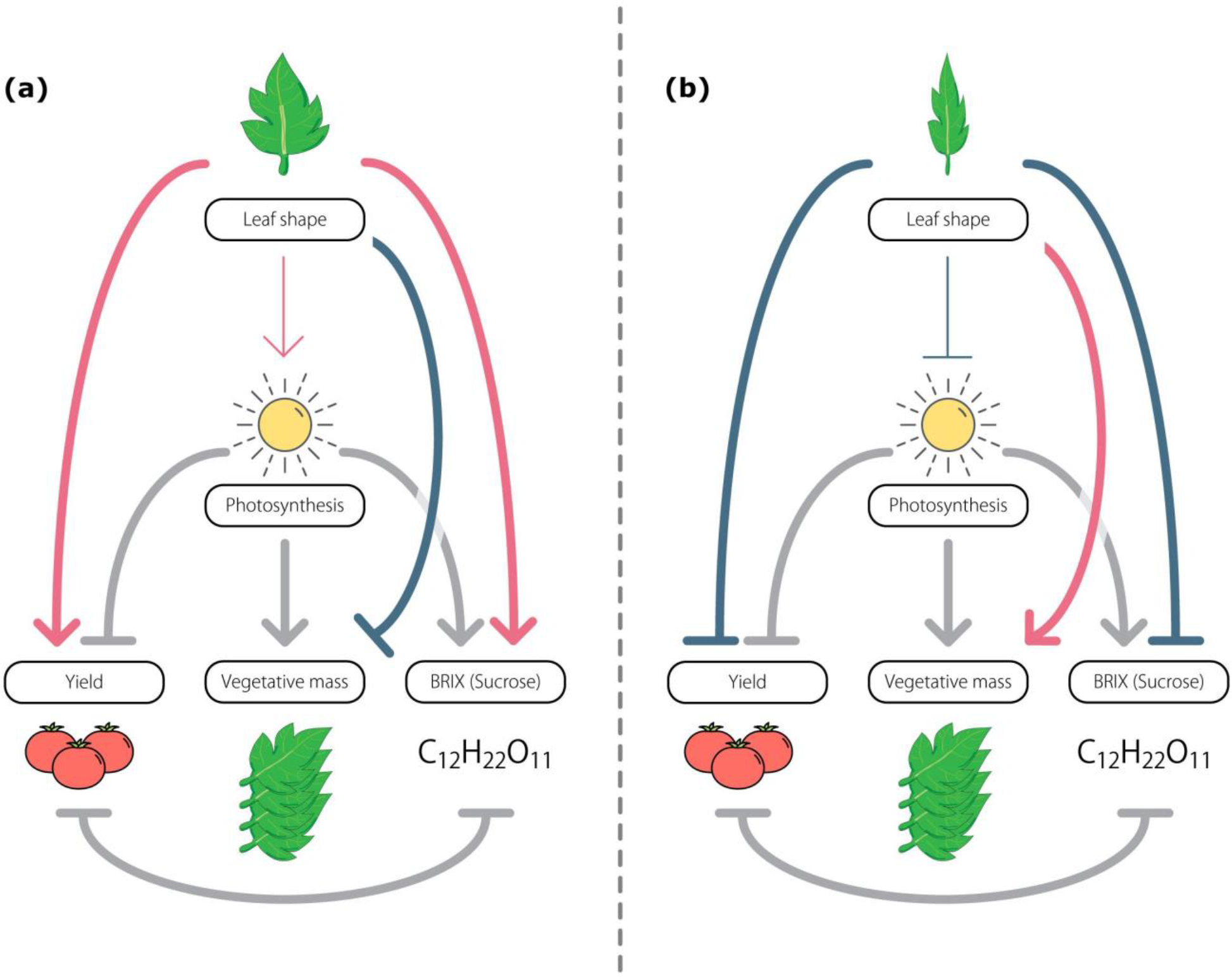
Composite model for leaf shape effects on fruit quality. The model was derived from the PLS-PM analysis. (a) The effects of round leaves on fruit quality and photosynthesis. (b) The effects of narrow leaves on fruit quality and photosynthesis. Red lines indicate a positive interaction while blue lines indicate a negative interaction. Grey lines indicate that the relationship does not change between the two leaf shapes. Narrower lines indicate a weaker relationship between the two connected traits.

We have shown that leaf shape strongly impacts the overall fruit quality in tomato, with rounder, less lobed leaves giving rise to higher yield and higher fruit BRIX. Photosynthesis, surprisingly, has a negative impact on yield while still positively contributing to fruit BRIX. Using data from cultivars not included in making our path model, we also showed that the model has a strong predictive performance for linking leaf shape to BY and could be used to potentially predict the outputs of a cultivar using leaf shape data (Table 2, 3). Our work shows the importance leaf shape to yield and BRIX across a wide array of genetic backgrounds, implicating leaf morphology in playing a significant and previously unidentified role in tomato fruit quality.

## Acknowledgements

We acknowledge help from Mary Lo Lee, Kirsten Brand, Gabriel Luis Moreira, Kristina Khuu, Eduardo Ramirez, Divya Kumaria, Jason Kao, and Surbhi Chopla in field sample collection, leaf and BRIX data collection, whole plant imaging, and seed collection. We also thank Diane Beckles, Donnelly West, and Jessica Budke for advice on data analysis. Support from the USDA-NIFA (grant number 2014-67013-21700) is gratefully acknowledged.

## Author Contribution

Neelima Sinha, Julin Maloof, and Dan Chitwood conceived of the research idea; Julin Maloof gave advice on research and analysis methods; Zizhang Cheng and Hokuto Nakayama did the whole genome sequencing; Amber Flores helped designing the sugar assays and collecting data; Kristina Zumstein organized the field study, helped with data collection, and all leaf analyses; Steven Rowland led the research effort, collected and analyzed data and wrote the paper with input from all authors; Neelima Sinha supervised the research and helped write the manuscript.

## Supporting Information

**Fig. S1** Yield and Fruit BRIX whole season measurements.

**Fig. S2** Whole plant adjusted *A* and *gst*.

**Fig. S3** Mean leaf complexity of all eighteen heirloom cultivars.

**Fig. S4** Alignment of sequence overhangs in the third exon of the *C*-Locus with the Rider TE sequence.

**Fig. S5** Mean shape from efourier and PCA analysis of eight heirloom cultivars.

**Fig. S6** PLS-PM inner and outer model representation and their correlations.

**Table S1** Outer model loadings and significance.

**Table S2** R^2^ values for each Latent Variable in the inner model.

**Table S3** Predictive performance and model fit for Heirloom cultivars in PLS-PM

