## Supplemental Information for "Leaf shape is a predictor of fruit quality and cultivar performance in tomato"

Article acceptance date: Click here to enter a date.

The following Supporting Information is available for this article:

**Fig. S1** Yield and Fruit BRIX whole season measurements.

**Fig. S2**	Whole plant adjusted *A* and *gst*.

**Fig. S3**	Mean leaf complexity of all eighteen heirloom cultivars.

**Fig. S4** Alignment of sequence overhangs in the third exon of the C-Locus with the Rider TE sequence.

**Fig. S5** Mean shape from efourier & PCA analysis of eight heirloom cultivars.

**Fig. S6** PLS-PM inner and outer model representation and their correlations.

**Table S1** Outer model loadings and significance.

**Table S2** R^2^ values for each Latent Variable in the inner model.

**Table S3** Predictive performance and model fit for Heirloom cultivars in PLS-PM.

**Fig. S1 Yield and Fruit BRIX whole season measurements.** a) The yield for each heirloom cultivar across fourteen weeks of the growing season is shown. Yield was calculated as kg of fruit per plant. Bloody Butcher and Stupice show the most increase in yield near the end of harvest weeks (17 – 23). b) Fruit BRIX was collected along with yield for each collection. The BRIX for most of the lines remained stable over the course of the season, between unripe and ripe fruit, with the exception of ABC Potato Leaf which increased by 2° BRIX over the last few weeks.

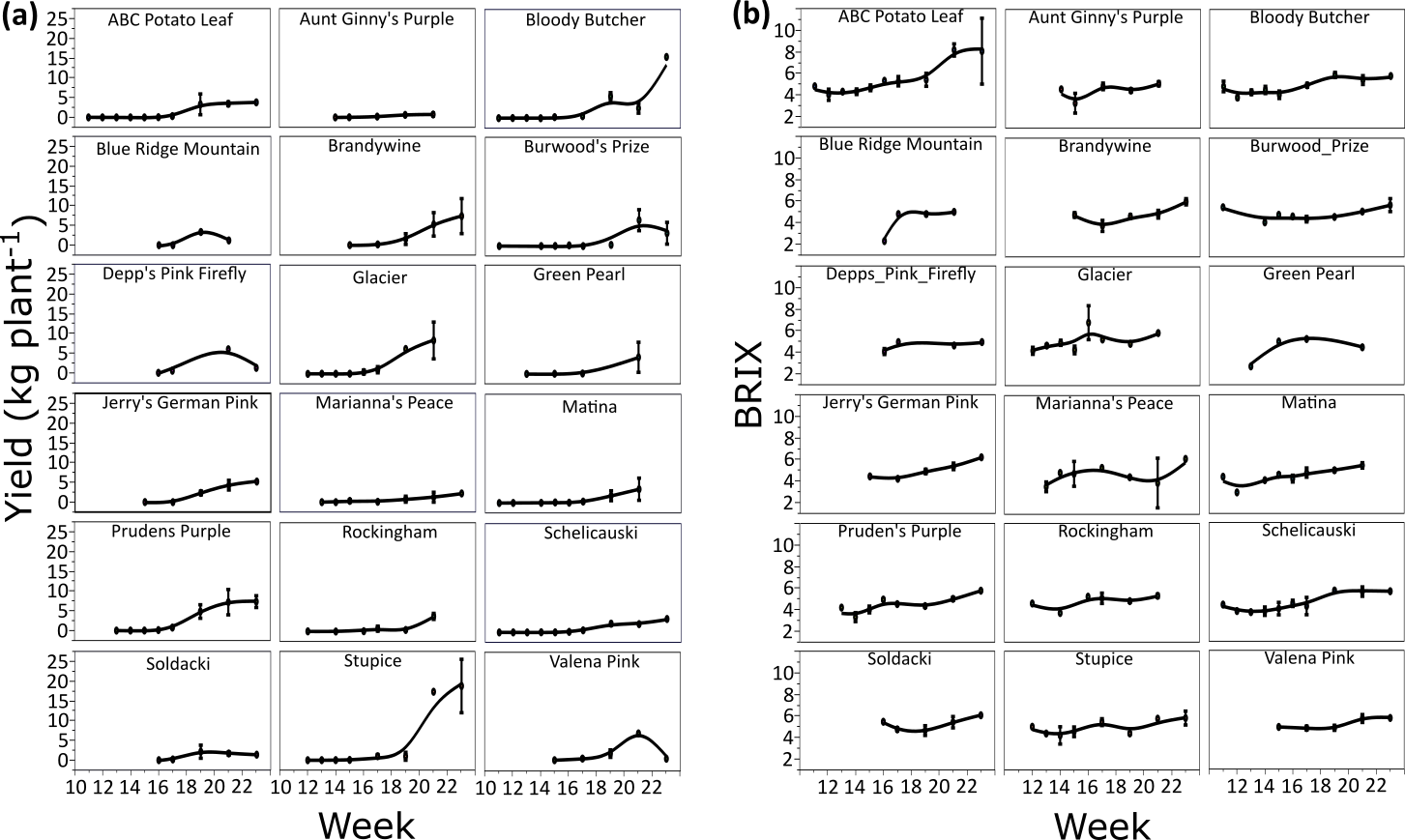

**Fig. S2**	**Whole plant adjusted *A* and *gst*.** To obtain a whole plant *A* rate and *gst* these values were multiplied by the total leaf area measured giving a per plant rate. The correlation between these values becomes nearly linear instead of logarithmic.

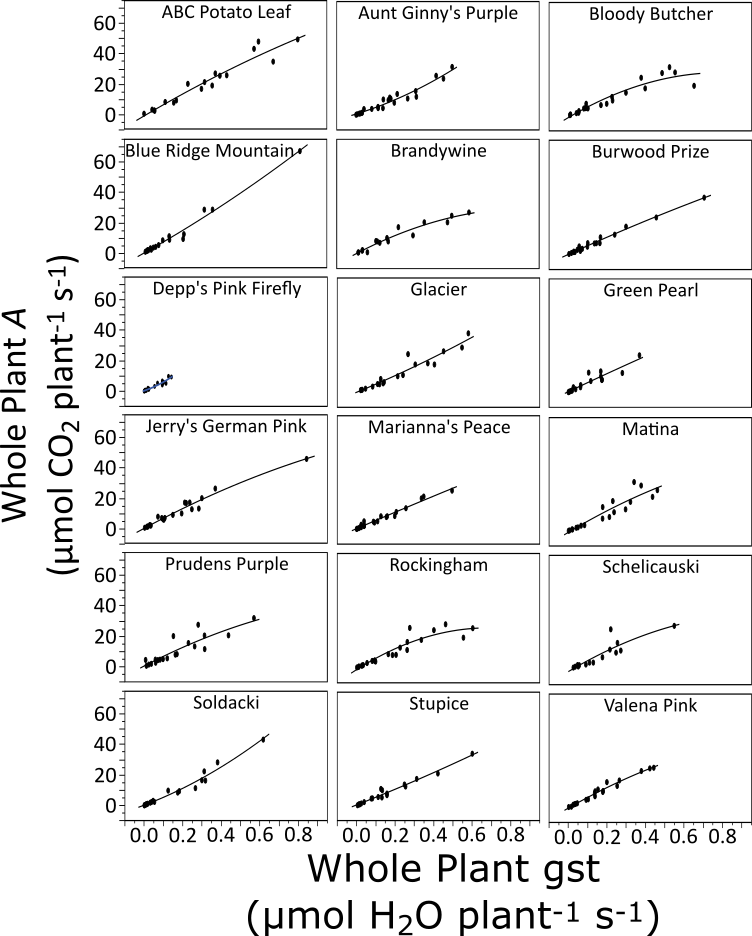

**Fig. S3**	**Mean leaf complexity of all eighteen heirloom cultivars.** The mean leaf complexity for all heirloom cultivars analyzed is shown. Glacier has the highest leaf complexity at 15, and Stupice the lowest at ~4. Error bars represent standard error.

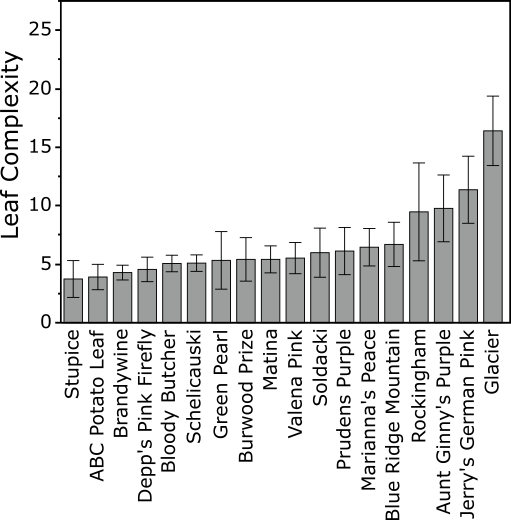

**Fig. S4**	**Alignment of sequence overhangs in the third exon of the C-Locus with the Rider TE sequence.** Six of the eight sequence heirloom cultivars contained overhangs in the same region of the third exon. These sequences were blasted and matched the CopiaSL_37/Rider sequence. The alignment of the first 100 base pairs and last 96 base pairs of the Rider element with the sequencing overhangs is shown. The middle portion of the Rider sequence has been removed as there was minimal mapping. The vertical dashed line shows the break between regions.

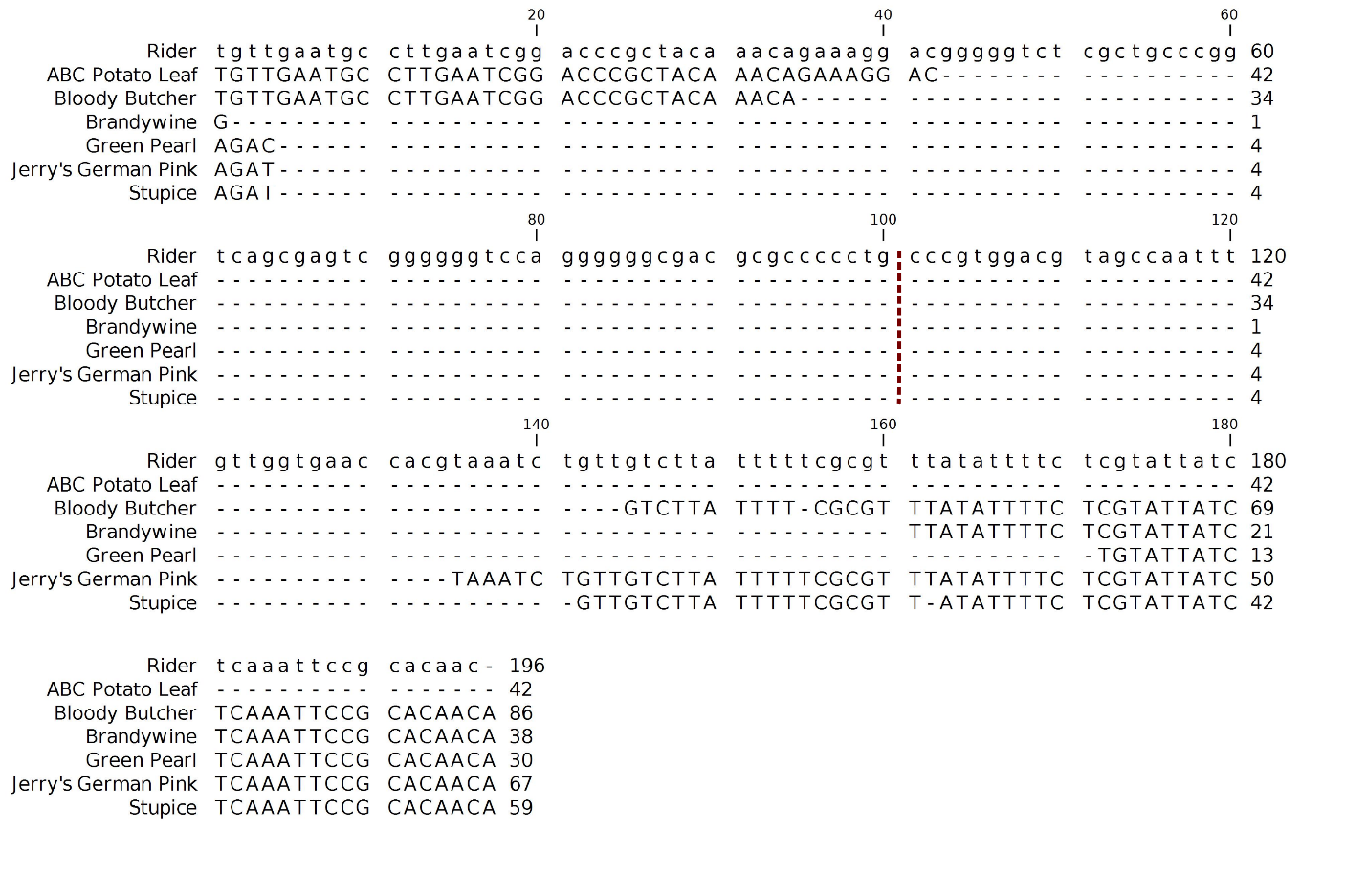

**Fig. S5** **Mean shape from efourier & PCA analysis of eight heirloom cultivars.** The mean shapes for each of eight heirloom cultivars is shown. These were generated using efourier analysis followed by PCA. These represent the relative shape and not size of the leaflets.

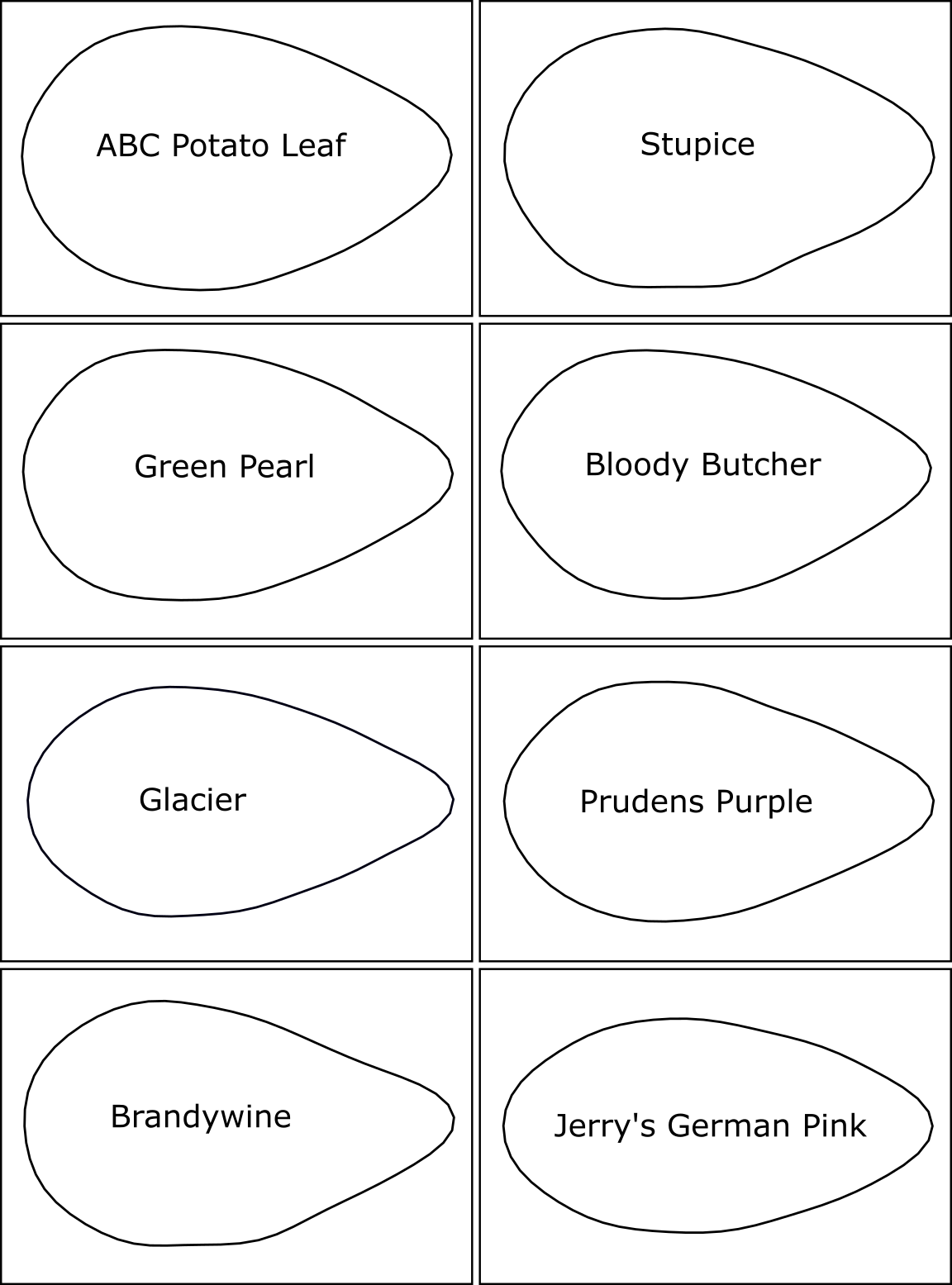

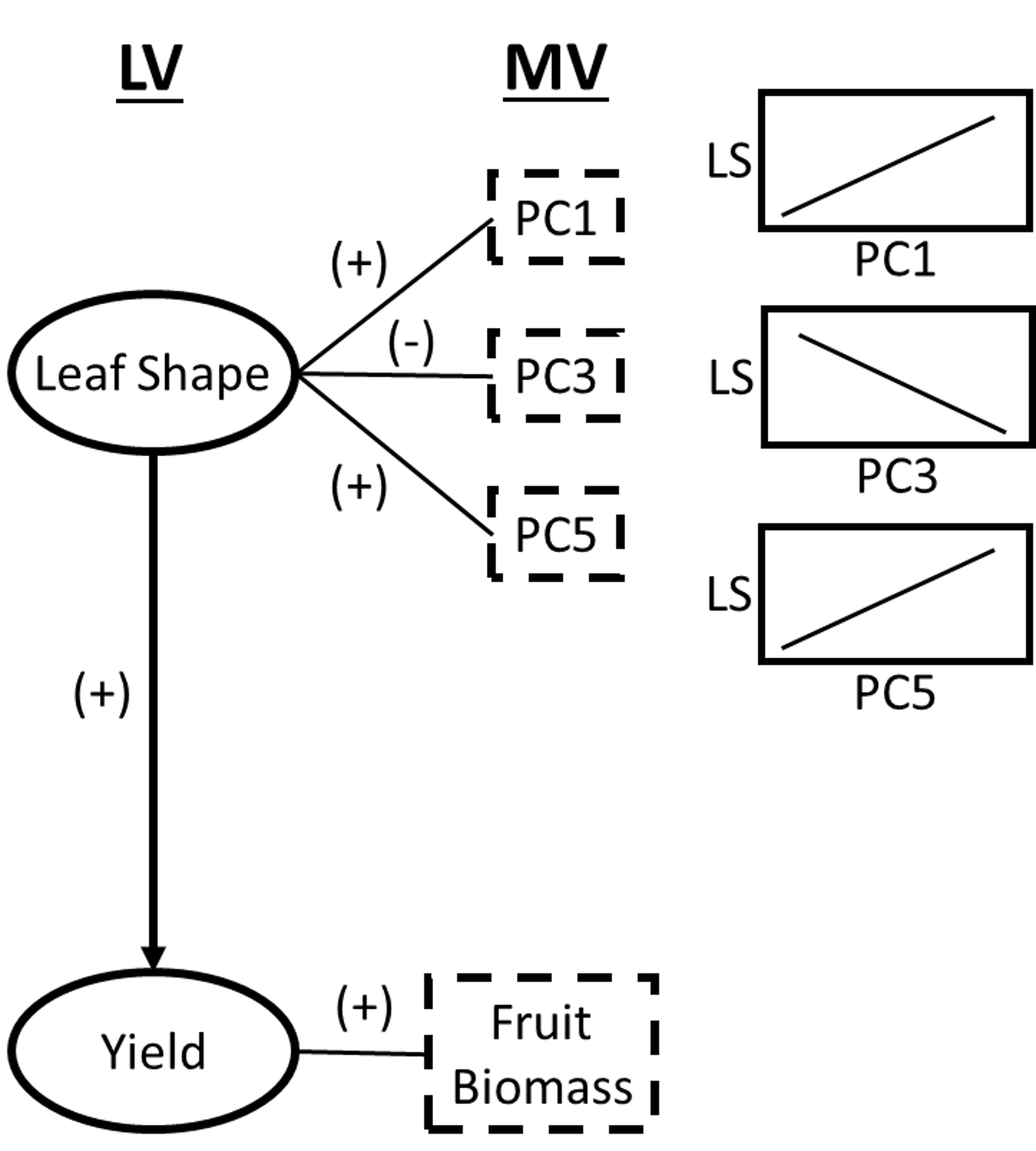
**Fig. S6** **PLS-PM inner and outer model representation and their correlations.** Manifest Variables (MV) are measured variables and are linearly correlated with their associated Latent Variable (LV), which is defined as the composite value of all it’s MVs. The causative relationship between LVs represents the modeled relationship between all associated MVs. As the value of Leaf Shape increases the value of Yield also increases. (+) represents a positive correlation and (-) a negative correlation.

**Table S1** Outer model loadings and significance. Original and sample mean for each manifest variable are shown, with standard deviation and P-values.

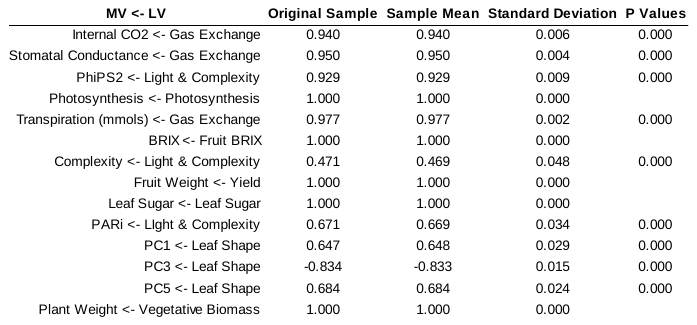

**Table S2** R^2^ values for each Latent Variable in the inner model. Original and sample mean values R^2^ values are shown.

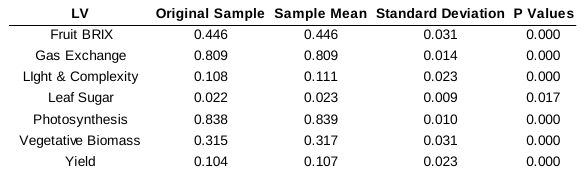

**Table S3** Predictive performance and model fit for Heirloom cultivars in PLS-PM. The RMSE, MAE, MAPE, and Q2 for the complete PLS-PM model, ABC Potato Leaf, and Aunt Ginny’s Purple is shown.

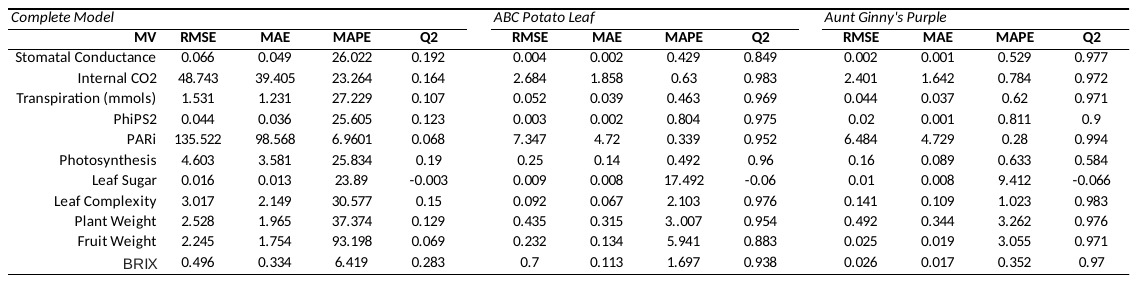

RMSE, Root Mean Square Error; MAE, Mean Absolute Error; MAPE, Mean Absolute Percent Error.
